# Effects of acute and repeated administration of the selective M_4_ PAM VU0152099 on cocaine vs. food choice in male rats

**DOI:** 10.1101/2021.09.06.459176

**Authors:** Morgane Thomsen, Jill R. Crittenden, Craig W. Lindsley, Ann M. Graybiel

## Abstract

Ligands that stimulate muscarinic acetylcholine receptors 1 and 4 (M_1_, M_4_) have shown promising effects as putative pharmacotherapy for cocaine use disorder in rodent assays. We have previously shown reductions in cocaine effects with acute M_4_ stimulation, as well as long-lasting, delayed, reductions in cocaine taking and cocaine seeking with combined M_1_/M_4_ receptor stimulation or with M_1_ stimulation alone. M_4_ stimulation opposes dopaminergic signaling acutely, but direct dopamine receptor antagonists have proved unhelpful in managing cocaine use disorder because they lose efficacy with long-term administration. It is therefore critical to determine whether M_4_ approaches themselves can remain effective with repeated or chronic dosing. We assessed the effects of repeated administration of the M_4_ positive allosteric modulator (PAM) VU0152099 in rats trained to choose between intravenous cocaine and a liquid food reinforcer, to obtain quantitative measurement of whether M_4_ stimulation could produce delayed and lasting reduction in cocaine taking. VU0152099 produced progressively augmenting suppression of cocaine choice and cocaine intake, but produced neither rebound nor lasting effects after treatment ended. To compare and contrast effects of M_1_ vs. M_4_ stimulation, we tested whether the M_4_ PAM VU0152100 suppressed cocaine self-administration in mice lacking CalDAG-GEFI signaling factor, required for M_1_-mediated suppression of cocaine self-administration. CalDAG-GEFI ablation had no effect on M_4_-mediated suppression of cocaine self-administration. These findings support the potential usefulness of M_4_ PAMs as pharmacotherapy to manage cocaine use disorder, alone or in combination with M_1_-selective ligands, and show that M_1_ and M_4_ stimulation modulate cocaine-taking behavior by distinct mechanisms.

## Introduction

Cocaine use disorder is becoming recognized as a public health problem of epidemic proportions. Despite relatively stable numbers of cocaine users, cocaine-related overdose deaths are rapidly increasing and now rival or exceed opioid overdose deaths in some American populations^1^. In the European Union, cocaine is the most widely used illicit drug^2^. There is currently no approved pharmacological treatment for cocaine use disorder, and psychosocial treatments are inadequate with high rates of relapse^1^.

Stimulation of muscarinic acetylcholine receptors continues to be tested as a pharmacological treatment in cocaine use disorder, but results of clinically available drugs are complicated by low efficacy and/or side effects. Clinical studies have so far relied on acetylcholinesterase inhibitors (AChEIs), which boost cholinergic signaling unselectively by slowing the degradation of acetylcholine. AChEIs can decrease cocaine self-administration in rats and monkeys, albeit typically with some decreases in food-maintained responding as well^3–5^. In clinical studies in cocaine-dependent subjects, various AChEIs decreased positive subjective effects of cocaine but most often failed to decrease drug use^6–9^ but see ^10^. The ability of AChEIs to decrease drug taking may be limited by side effects that preclude the use of higher (effective) doses and by their actions at several receptors that may counteract each other.

In order to isolate the potentially beneficial effects of cholinergic stimulation, we evaluated ligands that selectively activate specific muscarinic receptor subtypes. Several lines of evidence point to M_1_, M_4_, and M_5_ receptors as targets for cocaine use disorder treatment^11,12^. M_1_ and M_4_ are the principal muscarinic subtypes in striatal tissues^11^. In humans, M_4_ receptor expression (mRNA and protein) was reduced in putamen from patients with alcohol use disorder relative to controls, a finding that was mirrored in dorsolateral striatum of alcohol-exposed rats^13^. Although no difference in striatal M_4_ receptor mRNA levels was detected between brains from cocaine-addicted patients and controls^14^, a genetic M_4_ receptor variant has been associated with risk of cocaine use disorder^15^, supporting the notion that dysregulated M_4_ signaling could contribute to the development of cocaine use disorder. In rats, cocaine exposure has been reported to cause persistent decreases in striatal muscarinic receptor binding^16–18^, although which muscarinic receptor subtype(s) were affected was not determined and results varied depending on dosing regimen and abstinence duration^19^.

We previously showed that repeated administration of xanomeline, an M_1_/M_4_ receptor-preferring agonist, shifted behavior from cocaine-taking towards the alternative liquid food reinforcer in rats, with effects that persisted for a few days after ended treatment^20^. Moreover, in the same assay protocol, acute administration of an allosteric M_1_ receptor-selective partial agonist produced decreases in cocaine-taking for several weeks after administration^21^. Post-administration, lasting effects were also seen in an extinction and reinstatement procedure in mice in which combined stimulation of M_1_ and M_4_ receptors facilitated extinction of cocaine seeking, by either xanomeline or a combination of the allosteric M_1_ partial agonist VU0357017 and the M_4_ positive allosteric modulator (PAM) VU0152100, whereas neither VU0357017 nor VU0152100 alone had much effect^22^. On the other hand, M_4_ PAMs alone, as acute dosing, can reduce cocaine self-administration and diminish the discriminative stimulus effect of cocaine in mice^23,24^. Acute M_4_ PAM administration was also shown to reduce alcohol intake in rats^13^. Thus, M_4_ PAM consistently showed therapeutic promise in attenuating the effects of abused substances acutely. Studies evaluating repeated or subchronic administration regimens of M_4_ PAMs on cocaine-taking, however, have been lacking.

Here, we assessed whether M_4_ receptor stimulation alone was sufficient to modulate cocaine vs. food choice behavior in rats, and whether it can produce lasting suppression of cocaine taking, during and after treatment. To this end, we tested acute and repeated administration of the brain-penetrant M_4_-selective PAM VU0152099^25^. We previously have found that M_1_ receptor partial agonist suppression of cocaine self-administration was dependent on the signaling factor CalDAG-GEFI^26^. Therefore, we also tested whether M_4_ PAM-mediated suppression of cocaine self-administration is similarly dependent on CalDAG-GEFI signaling as a way to determine whether similar signaling pathways were affected by M_4_ and M_1_ receptors in the context of cocaine use disorder.

## Materials and Methods

### Subjects and housing

Male Sprague-Dawley rats were acquired at 7-8 weeks of age from Charles River (Wilmington, MA). Male and female mice lacking CalDAG-GEFI and wildtype sibling controls were generated by intercrossing heterozygous knockout mice in a congenic C57BL/6J genetic background at the Massachusetts Institute of Technology, as previously described^26^. Animals were allowed to acclimate to the laboratory for at least a week before training began. Rats were group-housed, up to 4 animals per cage until surgery, then individually housed post-operatively. Mice remained group-housed throughout. Rats had free access to water and were fed ~17g standard rodent chow daily (Diet 5001; PMI Feeds, Inc., St. Louis, MO) to maintain a 400-550 g bodyweight. Mice had access to food and water ad libitum. For enrichment, species-appropriate treats were provided once or twice weekly (Bio-Serv, Frenchtown, NJ), as were nesting and hiding materials. Mice had exercise devices before catheter implantation only, to avoid potential injury caused by the protruding catheter base post-surgery. Facilities were maintained on a 12-h light/dark cycle and testing was conducted during the light phase. Husbandry and testing complied with the guidelines of the National Institutes of Health Committee on Laboratory Animal Resources and the EU directive 2010/63/EU. All protocols were approved by the McLean Hospital Institutional Animal Care and Use Committee.

### Apparatus

Modular rat and mouse operant conditioning chambers and associated hardware from MED Associates Inc. (Georgia, VT) were used, each placed within a sound-attenuating cubicle equipped with a house light, a fan, and one (mice) or two (rats) syringe pumps (3.3 rpm, MED Associates model PHM-100) for the delivery of liquid food and i.v. cocaine, respectively, through Tygon tubing. Cocaine was delivered using a single channel fluid swivel (rats: MS-1, Lomir Biomedical, Malone, NY; mice: 375/25; Instech Laboratories, Plymouth Meeting, PA) mounted on a balance arm, which allowed animals free movement. In rat chambers, manipulanda were three retractable response levers (model ENV-112CM), two “reinforcer” levers (referred to as the “left” and “right” levers) on one wall and a third “observer” lever centered on the opposite wall. A steel cup between the reinforcer levers served as a receptacle for the delivery and consumption of liquid food reinforcers. A three-light array, red, yellow, and green (ENV-222M), located above the right lever was illuminated to signify the availability of food, an identical array with one additional yellow light, above the left lever, was used to signal the cocaine dose available. A white light (ENV-229M) was located above the observer lever. In mouse chambers, manipulanda were two illuminated nose-poke holes (ENV-313M).

### Catheter implantation and maintenance

Animals were implanted with chronic indwelling i.v. catheters (Camcaths, Cambridge, UK) under isoflurane (rats) or sevoflurane (mice) vapor anesthesia, with catheters exiting at the midscapular region. Analgesic (ketoprofen 5 mg/kg) and antibiotic (amikacin 10 mg/kg) were administered perioperatively. Animals were allowed at least 7 days recovery before being given access to i.v. cocaine. During this period, a prophylactic dose of cefazolin (30-40 mg/kg) was delivered daily through the catheter. Thereafter, catheters were flushed daily with sterile saline containing heparin (3 USP U/0.1 ml). Catheter patency was verified by prominent signs of sedation within 3 seconds of infusion of a ketamine-midazolam mixture (15 + 0.75 mg/ml) through the catheter, and only data collected with demonstrated patent catheters were used.

### Operant conditions

Details of the rat choice assay have been described previously^20,27^. Rats were trained/tested in daily sessions Monday-Friday. Under the terminal schedule of reinforcement, daily sessions consisted of five 20-min task components separated by 2-min timeout periods. Responding was reinforced under a FR 1 concurrent FR 5 FR 5 chain schedule of reinforcement: responses on the right lever were reinforced with liquid food (75 μl of 32% vanilla flavor Ensure^®^ nutrition drink in water, Abbott Laboratories, Abbott, IL). Responding on the left led intravenous cocaine infusions of increasing dose for each task component: 0, 0.06, 0.18, 0.56, 1.0 mg/kg/infusion. Cocaine doses were achieved by varying the infusion time, adjusted individually according to bodyweight. Each component of a session started with one response-independent automated delivery of each reinforcer (cocaine, then food) at the dose that would be available in the component, and the illumination of the observer lever’s cue light to signify that responses had scheduled consequences. One response on the observer lever (i.e., the initial response lever in the chain schedule) caused the retraction of the observer lever, turned off the associated cue light, extended the left and right levers, and turned on the cue lights associated with the left and right levers to signify reinforcer availability. Specifically, illumination of a triple cue light above the right lever signaled food availability, and the light array over the left lever signaled the cocaine unit dose available: no light for 0, green for 0.06, green+yellow for 0.18, green+yellow+red for 0.56, and green+yellow+red+yellow for 1.0 mg/kg/infusion. When a reinforcer was earned, left and right levers retracted and their associated cue lights were turned off. After a 20-second timeout period (including the infusion time), during which responses had no scheduled consequences, the observer lever extended and its associated cue light was illuminated, starting a new trial. Per component, 15 total reinforcers were available (completion of the response requirement on the left lever during availability of the zero cocaine dose counted as one reinforcer). Choice training continued until behavior stabilized: three consecutive sessions with ≥5 reinforcers/component earned in components 1-4 and ≥1 reinforcer earned in component 5, and with the dose of cocaine producing >80% cocaine choice on any given day remaining within one-half log unit of the prior 3-day mean.

Once training was completed, we tested the effects of acute administration of VU0152099 (vehicle, 0.32, 1.0, 1.8, 3.2, and 5.6 mg/kg intraperitoneally) 30 min before session start. After each dose, rats again had to meet the criteria for stable baseline behavior in order to test again, with at least three sessions between doses. Based on the effects of acute dosing, we then selected 1.8 mg/kg/day as the dose of VU0152099 for evaluation of repeated (subchronic) treatment effects. VU0152099 was administered once daily for 7 consecutive days, starting on a Friday. Rats were injected but not tested on Saturday and Sunday (treatment day 2 and 3), while choice sessions were conducted on days 1, and 4 through 7. A priori power analysis based on effects of xanomeline in the choice assay^20^ or CalDAG-GEFI genotype by M_1_ agonist interaction^26^, respectively, with a power of 0.8, indicated a required n=4-6 rats or mice.

Mouse self-administration was performed as previously described^26,28^. Mice were first assessed in food-reinforced (vanilla-flavor Ensure^®^) responding under a fixed-ratio 1 (FR1) timeout 20s schedule of reinforcement, followed by extinction, and food concentration-effect determination. After catheter implantation, mice were allowed to self-administer 1 mg/kg/infusion cocaine until baseline criteria were met (two consecutive sessions with at least 15 reinforcers earned per session and at least 70% of responses in the active hole). Thereafter saline was substituted until responding extinguished to <66% of baseline, then baseline behavior was re-established with 1.0 mg/kg/infusion, followed by dose-effect functions determined within-subjects (saline, 0.032, 0.1, 0.32, 1.0 and 3.2 mg/kg/infusion, tested according to a Latin-square design). The 0, 0.1, 0.32, 1.0 cocaine unit doses were then presented again with pretreatment of VU0152100 1 mg/kg IP, 30 min. before the session. WT and CalDAG-GEFI^-/-^ mice performed comparably in food reinforcement conditions (not shown) and cocaine-reinforced operant behaviors^26^ in the absence of pre-treatments (baseline behaviors).

### Data Analysis

For the cocaine vs. food choice assay, the primary dependent variables recorded for each component were: number of cocaine injections earned, number of food reinforcers earned, and percent cocaine choice, calculated as (number of reinforcers earned on the cocaine-associated lever ÷ total number of reinforcers earned) x 100. Total rate of responding (all three levers) and rate of responding on the reinforcer-selection levers alone were also recorded and analyzed. Total cocaine intake in mg/kg per session, total food reinforcer intake in ml per session were also calculated for each rat. The percents of cocaine choices were used to calculate A_50_ values (potency), defined as the dose of cocaine that produced 50% cocaine choice in each rat, and determined by interpolation from two adjacent points spanning 50% cocaine choice. In instances in which cocaine choice was >50% at the lowest cocaine dose, a value of 0.032 mg/kg/injection was used as a conservative estimate for inclusion in statistical analyses (i.e., quarter-log below the lowest cocaine dose tested, 29% of values). In instances in which where cocaine choice was <50% at the highest cocaine dose, a value of 1.78 mg/kg/injection was used (quarter-log above the highest cocaine dose tested, 4% of values). Group means and standard errors of the means were calculated from the log(10) of individual A_50_ values.

Effects of VU0152099 pretreatment on percent cocaine choice, cocaine reinforcers, liquid food reinforcers, and rates of responding per component were analyzed by repeated-measures ANOVA with Greenhouse-Geisser correction (or if values were missing, mixed effects analysis) with the factors cocaine dose and treatment day (baseline vs. treatment for acute dosing experiments; baseline, day 1, days 4 through 7 for the repeated dosing experiment). Significant effects of treatment, or of treatment by cocaine dose interactions, were estimated post-hoc by Sidak multiple comparisons tests vs. baseline. VU0152099 acute doses could not all be tested in each rat (within-subject), therefore each VU0152099 dose was analyzed separately to allow for within-subjects analysis of treatment vs. baseline. For mouse self-administration, cocaine reinforcers were similarly analyzed by ANOVA with cocaine dose and pretreatment as repeated-measures variables and genotype and between-subjects variable, followed by two-way analysis with genotypes combined.

Pilot studies using cocaine discrimination in mice (data not shown) suggested that M_4_ receptor stimulation modulates behavioral effects of cocaine following a biphasic function, presumably due to some competing/opposing effects. Total cocaine intake and total liquid food intake per session, and log-transformed A_50_ values, were therefore each compared by mixed-effects analysis with test for trend (linear and non-linear), with VU0152099 dose as repeated-measures factor in the acute dosing experiment, and session as repeated-measures factor in the repeated dosing experiment. Power analyses were performed using G*power 2, StataSE v.13 was used for 3-way ANOVA, and Graphpad Prism version 8 was used for all other analyses; *p*<0.05 is described as significant.

### Drugs

Cocaine hydrochloride was provided by the National Institute on Drug Abuse, National Institutes of Health (Bethesda, MD), dissolved in sterile 0.9% saline, and kept refrigerated. VU0152099 and VU0152100 (no salt) were synthesized at Vanderbilt University as previously reported (Brady et al. 2008). VU0152099 and VU0152100 were prepared daily by stirring in lukewarm Tween80 and diluted with sterile deionized water to the desired dose and to 5% Tween80.

## Results

### Repeated dosing choice studies

We traced the evolution of the effects on choices to obtain cocaine vs. a liquid food reinforcer over repeated days of administrations of the M_4_ PAM, VU0152099. At baseline, cocaine vs. liquid food choice behavior was consistent with that previously reported in this assay, with rats choosing mostly food during the first two components (no cocaine or 0.06 mg/kg/infusion cocaine available), and mostly cocaine during the last three components (0.18-1.0 mg/kg/infusion cocaine available). Fig. 1 shows percent cocaine choice, cocaine reinforcers, and liquid food reinforcers taken as functions of cocaine unit dose at baseline and on days 1 and 7 of repeated daily administration of 1.8 mg/kg VU0152099 (Fig. S1 shows all test days and rates of responding). Statistical analysis confirmed that cocaine self-administration and percent cocaine choice decreased over the week of treatment, reflected by significant effects of treatment day ([F(3,23)=3.6,*p*=0.02], [F(2,16)=4.1, *p*=0.03], respectively), and, for cocaine reinforcers, a cocaine by day interaction [F(20,125)=1.9, *p*=0.01] (full statistical analysis is reported in Table S2A). Food reinforcers taken and rates of responding were not affected systematically. Cocaine choice and cocaine and food reinforcers were related to cocaine unit dose (p≤0.003, Table S2A). Post-hoc comparisons relative to baseline reached significance on day 7, at the 0.18 mg/kg cocaine unit dose, for both percent cocaine choice (p=0.02) and cocaine reinforcers (p=0.02). Of the six rats tested for all seven treatment days, all but one showed a shift in cocaine choice by day 7;Fig. S3 shows choice curves in individual rats.

**Figure 1.**
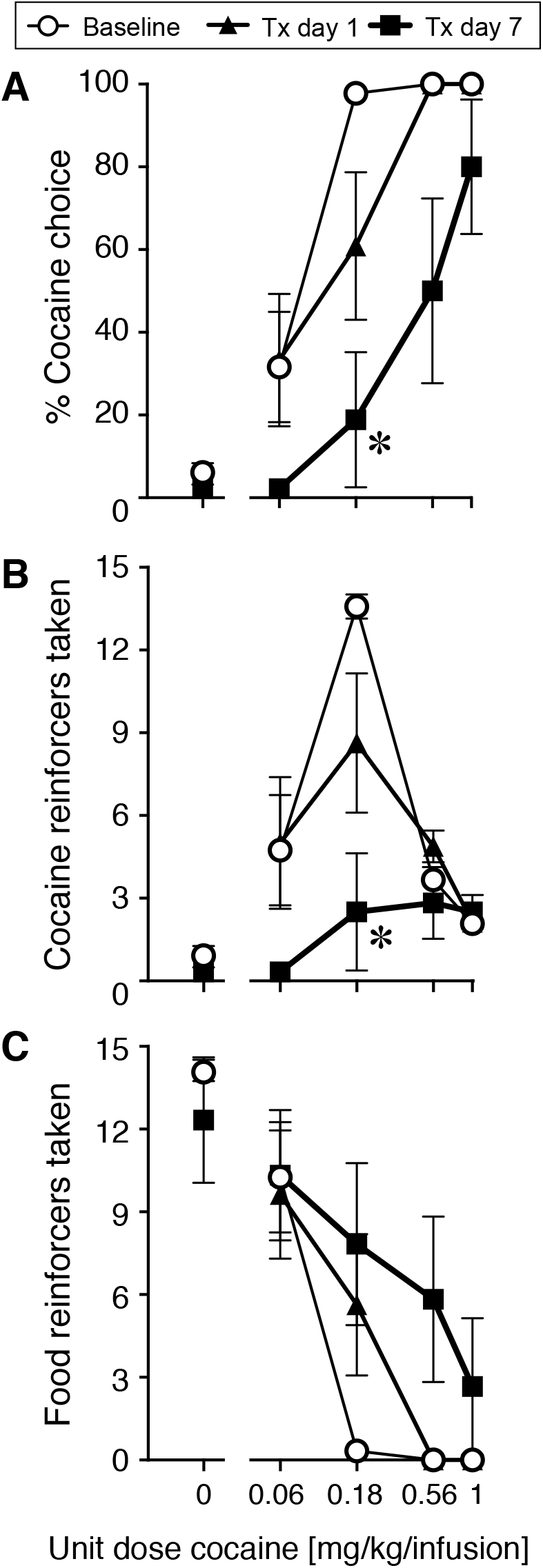
Cocaine vs. food choice behavior in rats during acute (day 1) and repeated (7 days) pretreatment with 1.8 mg/kg VU0152099. Allocation of behavior (A) shifted away from cocaine towards food taking during treatment. Cocaine self-administration (B) was markedly suppressed after a week of daily treatment, while food taking (C) was increased. *p<0.05 vs. baseline.

Gradual shifts in the choice curve, from cocaine taking towards food taking, were indicated by a near-significant effect of treatment day on A_50_ values [F(3,15)=3.4, p=0.054] and a significant linear trend [F(1,38)= 11.3, p=0.002] (Fig. 2A, Table S2B). There was similarly a significant linear trend for decreasing session-total cocaine intake [F(1,32)=9.7, p=0.004] (Fig. 2B).

**Figure 2.**
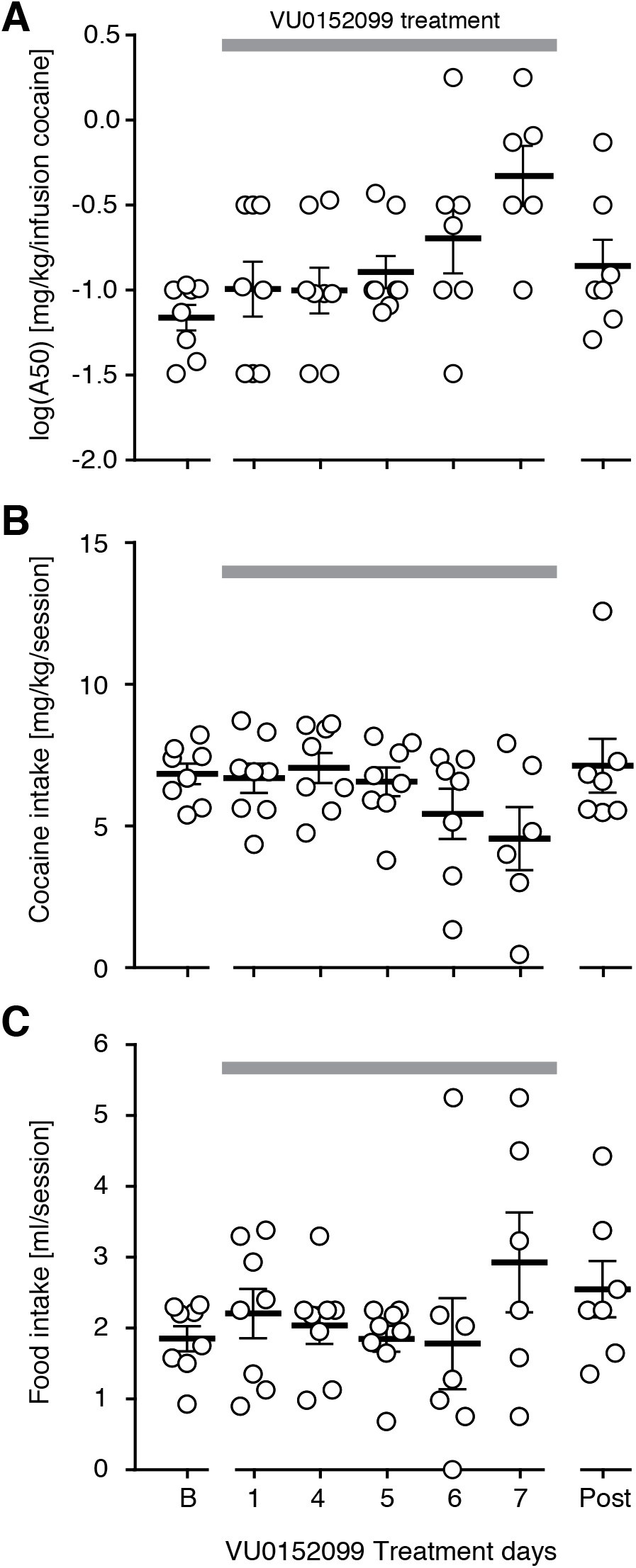
Effects of daily VU0152099 treatment in the rat cocaine vs. food choice over days, grey bar indicates days of VU0152099 administration. (A) The potency of cocaine in the choice assay, calculated as the dose of cocaine to maintain 50% choice (A50) showed a linear trend for more cocaine needed to maintain cocaine choice. (B) Session-total cocaine intake showed a linear decreasing trend over treatment time. (C) Food intake was not affected systematically, with only non-significant tendency to increase over treatment time. “B”: Baseline, “post”: post-treatment, day four after treatment ended.

No consistent change in cocaine or food intake was observed in the two rats tested with repeated vehicle administration, the only noticeable changes from baseline behavior being an increase in cocaine intake after the weekend in one rat, and an increase in food intake after the weekend in the other rat - transient “Monday effects” that are not unusual also in rats maintained on baseline conditions (Fig. S4).

### Acute administration choice studies

The dose used for the repeated-dosing study was selected based on the acute effects of a range of VU0152099 doses. Fig. 3 shows effects of acute pretreatment with VU0152099 on percent cocaine choice, cocaine reinforcers taken, and food reinforcers taken as functions of cocaine unit dose. All doses of VU0152099 appeared to cause a modest downward shift in the cocaine self-administration dose-response curve, with 1.8 mg/kg VU0152099 reaching statistical significance relative to baseline [F(1,6)=8.5, *p*=0.03]. Pairwise comparisons at each cocaine dose did not reach statistical significance. Food reinforcer curves showed non-significant trends for more food taken. Percent cocaine choice similarly showed consistent trends for a shift from cocaine to food taking (except at the highest dose of VU0152099), approaching significance for the 1.8 mg/kg VU0152099 dose (*p*=0.07 vs. baseline). Percent cocaine choice, cocaine reinforcers taken, and food reinforcers taken were again always significantly related to cocaine unit dose (see Table S2C for full statistical details). All rats returned to their baseline choice and intake levels within a few days of testing (data not shown), though not necessarily on the first day after testing (grey curves in Fig. 3).

**Figure 3.**
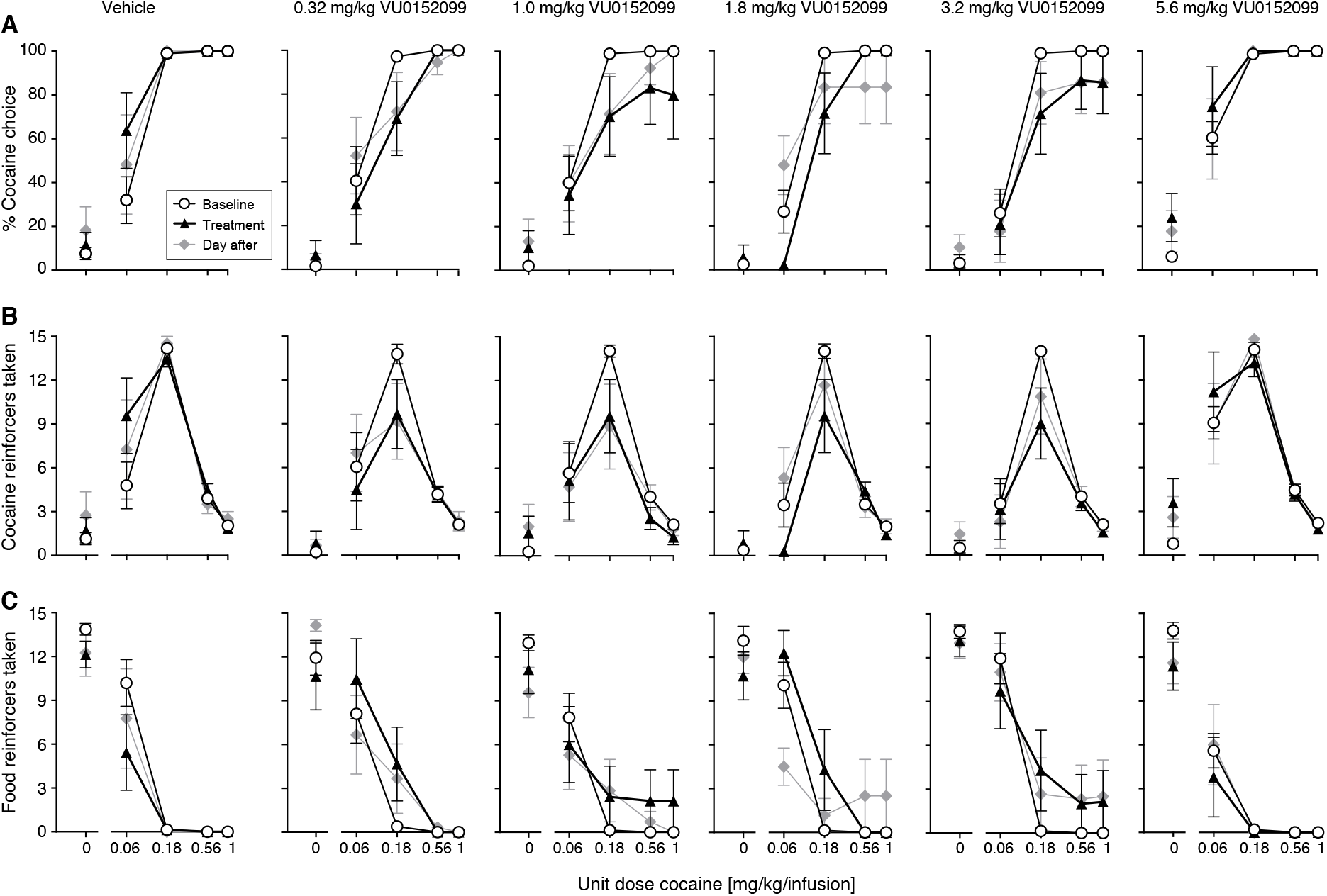
Cocaine vs. food choice behavior in rats after various acute doses of VU0152099. Cocaine vs. food choice allocation (A), cocaine self-administration (B), and food taking (C) showed modest effects of acute VU0152099 administration (black triangles) relative to baseline (open circles), with a significant main effect of treatment on cocaine self-administration for only 1.8 mg/kg VU0152099. There were no significant “day after” effects, baseline test after VU0152099 administration, showed in grey diamonds. Vehicle alone (leftmost panels) caused a momentary decrease in food taking.

Vehicle (5% Tween80 in water) treatment in itself affected cocaine vs. food taking behavior, relative to baseline, although in the opposite direction of VU0152099 (see Fig. 3 leftmost panels). Specifically, self-administration of the lowest cocaine dose (0.06 mg/kg/infusion) was increased (treatment effect [F(1,6)=6.7, *p*=0.04], cocaine by treatment interaction [F(2,9)=7.5, *p*=0.02]). Food reinforcers taken in the same component were decreased (treatment [F(1,6)=9.1, *p*=0.02], interaction [F(1,8)=7.6, *p*=0.02]). There was a significant treatment by cocaine dose interaction on percent cocaine choice [F(1,7)=7.9,*p*=0.02]. Vehicle administration did not affect rates of responding.

Total rates of responding were significantly related to cocaine unit dose, with slower rates as cocaine dose was increased, while rates of responding on the reinforcer-selection levers remained comparable across cocaine doses (see Fig. S5, and Table S2C). Conversely, the total rates were not affected by VU0152099 administration, but 0.32 mg/kg (*p*=0.001) and 5.6 mg/kg (*p*=0.02) VU0152099 increased rates of responding on the reinforcer-selection levers relative to baseline (Fig. S5).

Shifts in the choice curve as represented by change in the cocaine dose that produced 50% cocaine choice, relative to baseline, showed a biphasic curve with increased A_50_ up to 3.2 mg/kg VU0152099 (see Fig. S6A). The effect of VU0152099 dose approached significance (*p*=0.06, full statistical details in Table S2C), and showed a significant test for trend, nonlinear [F(5,32)=3.8, *p*=0.008]. The total cocaine intake per session similarly showed a significant nonlinear trend [F(5,33)=3.0,*p*=0.02] with apparent decrease in intake at intermediate doses, while liquid food intake showed a nonlinear trend [F(5,33)=2.8, *p*=0.03] with higher intake at intermediate doses of VU0152099 (Fig. S6B,C). These analyses indicate a dose-dependent, though modest, reallocation of behavior from cocaine-taking towards food-taking behavior.

### CalDAG-GEFI knockout mouse studies

We previously have found that CalDAG-GEFI mediates M_1_ dependent synaptic plasticity in striatal medium spiny neurons (MSNs)^26^. In self-administration tests of mice lacking CalDAG-GEFI, we found that the CalDAG-GEFI^-/-^ mice were completely insensitive to M_1_ partial agonist pretreatment that successfully eliminated cocaine self-administration in sibling control mice^26^. In order to test whether this effect generalized to M_4_ signalling, we therefore tested the M_4_ PAM VU052100 in cohorts of CalDAG-GEFI^-/-^ and WT mice that self-administered cocaine. VU052100 suppressed cocaine self-administration in all mice regardless of genotype (Fig. 4; effect of pretreatment [F(1,42)=19.4, p=0.0001], pretreatment by gene or pretreatment by gene by cocaine interactions p=0.6, see full analysis in Table S2D). Thus, the M_4_ PAM effect was not dependent on CalDAG-GEFI, differentiating further M_4_ and M_1_ functions in potential mediation of cocaine self-administration.

**Figure 4.**
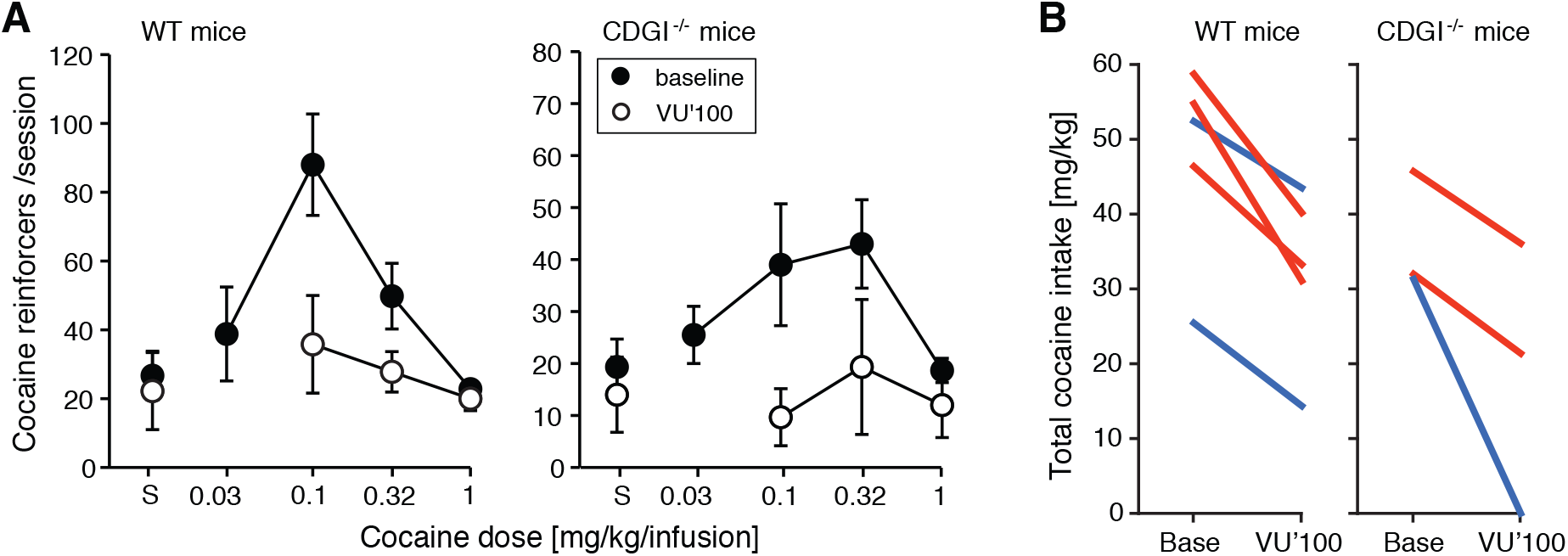
Effects of M_4_ PAM VU0152100 in WT and CalDAG-GEFI^-/-^ mice (CDGI^-/-^). (A) Group means of cocaine self-administration dose-effect functions with and without 1.0 mg/kg VU0152100 (VU’100) pretreatment. Data are cocaine reinforcers taken per 3h-session, n=5 WT (3 females, 2 males) and 3 CDGI^-/-^ (2 females, 1 male); “S”: saline. (B) Change in total cocaine intake as mg/kg (summed over doses) between baseline and VU0152100 treatment, showing all individual mice with females in red, males in blue.

## Discussion

Our findings demonstrate the reduction of cocaine self-administration by selective M_4_ PAM administration, and further indicate that this effect does not require M_1_ activity signaled through a guanine nuclear exchange factor action required for reduction of cocaine self-administration by M_1_ partial agonist treatment. To evaluate the effects of the M_4_ PAM, we used a paradigm requiring active choice between cocaine and a liquid food reinforcer. The advantages of choice assays over single-reinforcer assays are considerable^29,30^, but we note here as especially relevant the assessment of choice allocation rather than rates of responding. VU0152099 has been found not to affect motor function measured by time on the rotarod in rats, tested up to 100 mg/kg^25^. However, decreased food-reinforced operant behavior in mice treated repeatedly with the M_4_ PAM VU0467154 has been observed in tasks with higher complexity or response requirement (Justinussen et al., manuscript in preparation). Thus, M_4_ receptor stimulation could decrease motivated behaviors generally rather than suppress cocaine taking specifically, at least under some conditions, reducing clinical usefulness of M_4_ PAM treatments. Here, we found that treatment with VU0152099 evoked reallocation of behavior from cocaine taking towards a competing natural reinforcer with no suppression of overall rates of responding, even with repeated treatment. This result suggests preferential or selective suppression of cocaine-taking behavior by VU0152099, rather than general suppression. Conversely, a shift in cocaine vs. food choice can be achieved by modifying the reinforcing effects of cocaine or of the food^31^. We did not evaluate repeated M_4_ PAM administration in single-reinforcer assays in rats, but we did show M_4_-mediated suppression of cocaine self-administration in mice (see also Dall et al.^24^). In mice, M_4_ PAMs either had no effect on food-maintained responding or decreased it^24,32^, whereas an M_4_-preferring *antagonist* increased food-reinforced responding^33^. Thus, the observed shift in behavior allocation is most likely driven by reduced reinforcing (and/or discriminative stimulus) effects of cocaine, with a resulting increased *relative* value of the competing reinforcer, and not by increased intrinsic value of the food.

Another main objective of this study was to determine whether M_4_ receptor stimulation could maintain a suppressing effect on cocaine-taking behavior as repeated administration. A well-established effect of M_4_ receptor stimulation is functional dopamine antagonism, both via suppressed striatal dopamine release (electrically evoked or cocaine- or amphetamine-stimulated)^23,34,35^ and via opposing cellular effects of dopamine D_1_ receptor stimulation in MSNs^36,37^. Direct dopamine receptor antagonists have been evaluated both in laboratory animals (including in cocaine vs. food choice) and clinically, and uniformly decreased cocaine taking acutely but were found to be ineffective or increased cocaine taking and/or subjective effects of cocaine as chronic/repeated treatment^27^. Therefore, it was particularly important to assess M_4_ receptor stimulation over several days, as functional dopamine antagonism may be prone to the same problems. We found that VU0152099 not only shifted behavior away from cocaine taking during repeated treatment, but also that the effect grew larger over treatment days. VU0152099 has a half-life of 1.25 h in rat brain^25^, making accumulation in the blood or brain unlikely. In previous studies, removal of the cocaine reinforcer shifted behavior away from the cocaine-lever choice from the first day, but the calculated cocaine “intake” (i.e., mostly driven by reductions in the last component that previously produced the highest dose of cocaine) was reduced only from the second or third day (Fig. 5, ^31^). We hypothesize that VU0152099 decreases the reinforcing efficacy of cocaine, and that rats gradually extinguished cocaine-seeking in an extinction response. The difference between the effects of dopamine receptor antagonists and M_4_ PAMs is not necessarily surprising, since both suppression of dopamine release and suppression of dopamine receptor-induced second messenger cascades are complex effects and depend on stimulation conditions, rather than being universally dopamine-dampening^38,39^. Since PAMs amplify endogenous acetylcholine signaling, this complexity of the physiological modulation of dopaminergic signaling may be preserved, preventing the rapid development of tolerance seen with dopamine receptor antagonists.

**Figure 5.**
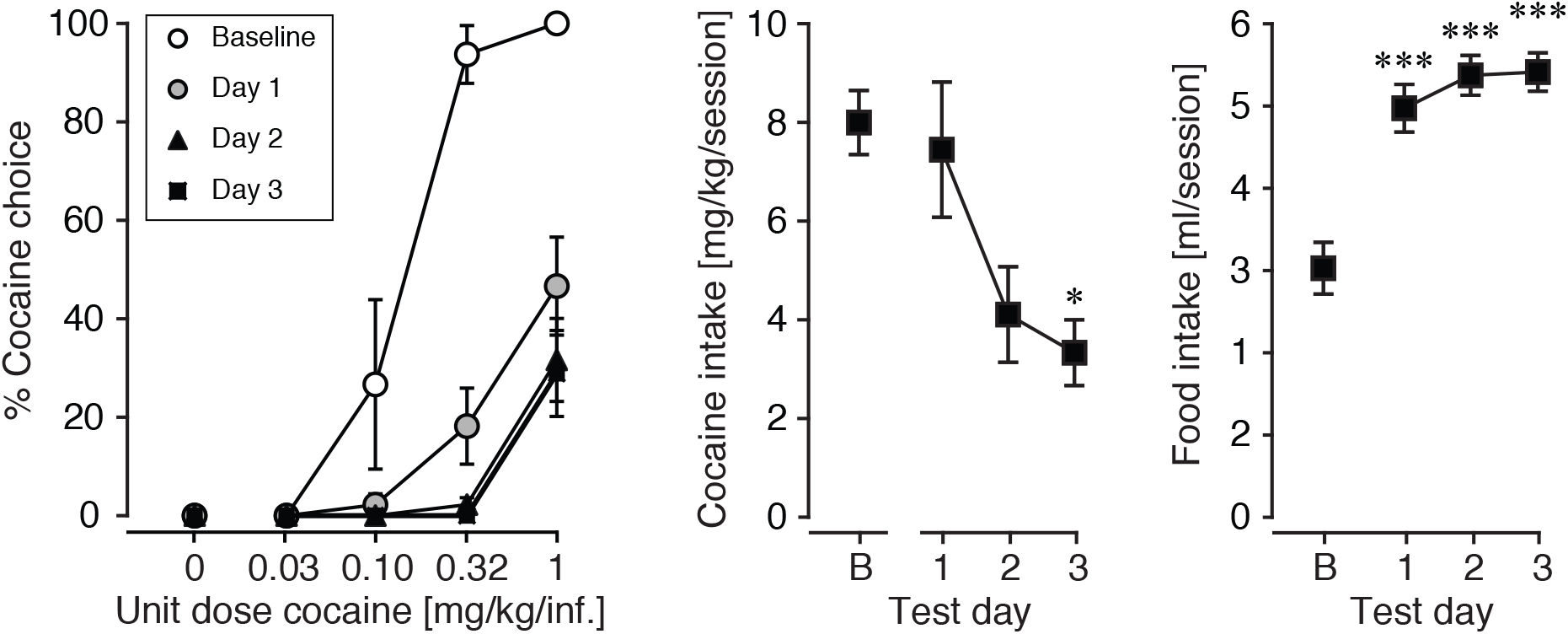
Effect of removing cocaine as the reinforcer in the rat choice assay. (A) Cocaine choice decreased when cocaine was removed for three consecutive sessions, all other contingencies remaining unchanged. (B) Session-wide cocaine intake did not decrease until the second day. (C) Increased allocation of behavior to the food-reinforced lever increased rapidly upon removal of cocaine. Data were previously published average over days (Thomsen et al 2013), and are reused with permission from John Wiley and Sons.

The effects of VU0152099 were maintained and grew over the treatment period, but rats returned to their pre-treatment levels of cocaine and food taking within days after the end of the treatment. This result stands in contrast to findings with M_1_ partial agonist treatment, which produced delayed and long-lasting (weeks) suppression of cocaine choice in this assay^21^. Xanomeline showed a similar but much briefer “post-treatment effect”^20^, and rats treated with the acetylcholinesterase inhibitors tacrine or donepezil showed decreased cocaine self-administration for several days after treatment ended^3,5^. Taken together with the present results, the findings suggest that the prolonged effects are attributable to action at M_1_ receptors, but not M_4_ receptors. The mechanisms mediating “anti-cocaine” effects of M_1_ and M_4_ stimulation are still not fully understood. Here, we were able to test one potential signaling cascade, based on our finding that the ability of M_1_ receptor partial agonist to suppress cocaine self-administration was dependent on the signaling factor CalDAG-GEFI^26^. Here we found that by contrast, the M_4_ PAM VU0152100 suppressed cocaine self-administration comparably in mice lacking CalDAG-GEFI and their control siblings, indicating that mechanisms different from M_1_ stimulation are involved. This difference accords with the fact that M_1_ and M_4_ receptors signal through different G-protein-coupled cascades. We have observed additive or synergistic effects of combined M_1_ and M_4_ receptor activation in at least some behavioral endpoints, such as facilitation of extinction, and studies using knockout mice indicate that both receptors mediate suppression cocaine’s discriminative stimulus effects^22,40^. A combination of selective M_1_ and M_4_ stimulation could thus provide both rapid-onset and long-lasting suppression of cocaine taking and cocaine choice, but this remains to be empirically determined.

M_4_ receptors are expressed in many brain regions including prefrontal cortex, hippocampus, thalamus, amygdala, and midbrain, but most densely in striatal tissues, where M_4_ receptors are found on cholinergic interneurons, on glutamatergic corticostriatal and thalamostriatal projections, and on (primarily D_1_ receptor-expressing) MSNs (for review, see ^11^). M_4_ receptors modulate striatal dopamine signaling but they also inhibit corticostriatal glutamatergic signaling onto both D_1_- and D_2_-expressing MSNs and modulate neuroplasticity^41,42^. These mechanisms could all contribute to reducing cocaine self-administration. Knockout mice lacking functional M_4_ receptors show a phenotype that could be interpreted as moderately “addiction-prone”. Specifically, M_4_ knockout (M_4_^-/-^) mice self-administered more cocaine and alcohol relative to wild-type controls and were slower to extinguish responses previously reinforced by cocaine or alcohol^43,44^. M_4_^-/-^ mice also emitted more responses when reinforced with liquid food^43^, suggesting the possibility of a more generally increased responsiveness to reinforcement. M_4_^-/-^ mice showed a mild compulsive-like and impulsive-like phenotype under some conditions in 5-choice serial reaction time tasks^45,46^ In mice lacking M_4_ receptors only on dopamine D1 receptor-expressing neurons, which are mostly MSNs (D1-specific M_4_^-/-^), VU0152100 still blunted cocaine-stimulated extracellular dopamine increase, but to a lesser degree than in intact mice^23^, suggesting that MSN and other M_4_ receptor populations are also involved. Dopaminergic signaling and behavioral hyperresponsiveness of global M_4_^-/-^ mice to cocaine was preserved in D_1_-specific M_4_^-/-^ mice^46,47^, but not in mice lacking M_4_ receptors on cholinergic neurons, which exhibited reward learning deficits^46^. Thus, M_4_ autoreceptors on cholinergic interneurons may not be a primary mediator of M_4_ “anti-cocaine” effects, M_4_ receptors on MSNs likely mediate some of the effects, leaving M_4_ receptors on glutamatergic projections as likely candidates for mediating a part of the effects. More studies are needed to clarify mechanisms of M_4_-mediated suppression of cocaine reinforcement.

Factors limiting the conclusions that can be drawn from the study include the evaluation of a single M_4_ PAM in the choice assay, and use of a different, but structurally related, M_4_ PAM in the mouse single-reinforcer assay. As pharmacological tools, VU0152099 and VU0152100 have shown high selectivity for M_4_ over other muscarinic receptor subtypes, and have shown clean ancillary pharmacology with only weak 5HT-2B receptor antagonist activity detected as off-target activity for VU0152099^25^. At the relatively low doses used in the present experiments, 5HT-2B receptor antagonism is unlikely to contribute to the effects of VU0152099, and systemic administration of 5HT-2B receptor antagonists has generally failed to modulate the behavioral effects of cocaine^48,49^. Nevertheless, the present results should be replicated using one or more well-characterized M_4_ PAMs, ideally structurally distinct. Vehicle treatment alone produced a small decrease in food taking early in the session, with a corresponding increase in cocaine taking at the lowest cocaine dose. Similarly, food taking was decreased in the first, no-cocaine component when VU0152099 doses were tested acutely, despite the shift from cocaine to food in later components, in which cocaine was available. These observations suggest that the Tween vehicle temporarily reduced food taking, perhaps due to nausea, but that these effects were either brief or overcome/masked by VU0152099 decreasing the reinforcing effect of cocaine. It is possible that shifts in cocaine vs. food choice would be larger, were they not confounded by vehicle effects. It also remains to be determined how well these findings translate across species, sex, age, degrees of cocaine exposure, and other parameters. Despite these caveats, out findings strongly indicate the pre-clinical value of M_4_ enhancement, and distinguish M_4_ signaling from that of M_1_ receptors in relation to the actions of CalDAG-GEF1 signaling: The repeated administration of VU0152099 produced behavioral shift in choice away from cocaine-taking toward a natural food reinforcer. The effect grew over treatment days, yet there was no prolonged or delayed “after effect”. This dynamics and the insensitivity of M_4_ PAM-suppression of cocaine self-administration to inactivation of the signaling factor CalDAG-GEFI demonstrate differences from M_1_-mediated suppression of cocaine reinforcement. We observed no adverse effects of acute or repeated administration of VU0152099 or VU0152100 in rats and mice, and no suppression of rates of responding in the choice assay. Nevertheless, development of M_4_ PAM for clinical use (primarily for the indication of schizophrenia) has been challenged by species differences in potency, selectivity over M_2_ receptors, CNS penetration, and in vivo efficacy^50^. M_4_ drug development continues to progress^50^, with at least one compound in early clinical trials (clinical trial identifier NCT04136873; NCT04787302).

## Acknowledgments

This research was supported by a grant from the National Institutes on Drug Abuse (DA027825, MT), and funding from the Saks-Kavanaugh Foundation (AMG) for the experiments with CalDAG-GEFI. While preparing the manuscript, MT was supported by funds from the Independent Research Fund Denmark (8020-00110B, MT). Discovery and development of VU0152099 and VU0152100 was supported by the Molecular Libraries Probe Production Centers Network (U54MH084659, CWL). The authors thank Christopher Adam, Kevin Stoll, and Benoit Niclou for technical assistance.

## Data sharing

The data that support the findings of this study are available from the corresponding author upon reasonable request.

## Authors contribution

MT conceptualized and designed the study, oversaw data collection, analyzed and interpreted the data, prepared figures. MT wrote the manuscript with input from JRC and editorial work by AMG. CWL synthesized and characterized VU0152099 and VU0152100. JRC and AMG generated the CalDAG-GEFI mice. All authors approved the final version for publication.

