## Supplemental Figure S1 for "Effects of acute and repeated administration of the selective M_4_ PAM VU0152099 on cocaine vs. food choice in male rats"

**Supplemental Figure S1.** Cocaine dose-response curves for all days, repeated VU0152099 dosing

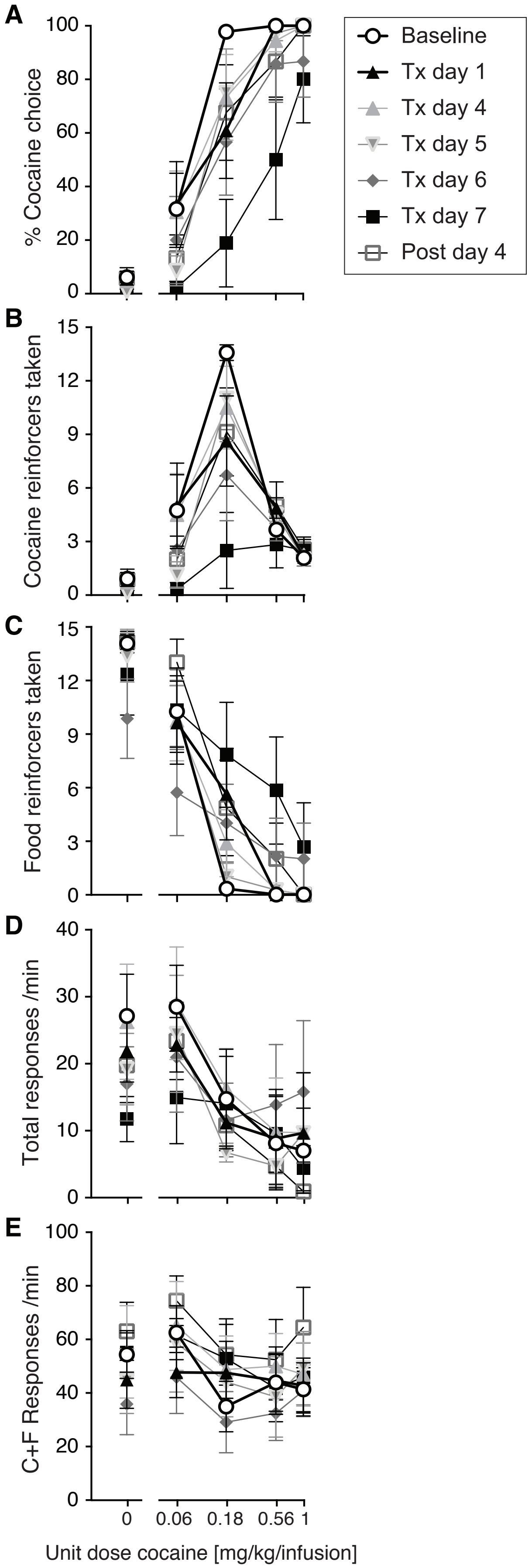


These are the same data as Fig. 1 but showing all test days, and rates of responding.
(A) % cocaine choice, (B) cocaine reinforcers taken, (C) liquid food reinforcers taken, (D) total rates of responding, (E) rate of responding on the two reinforcer-selection levers, as functions of cocaine unit dose during repeated daily administration of 1.8 mg/kg VU0152099.
Cocaine choice allocation and cocaine self-administration decreased over the week of treatment. Food reinforcers taken were not affected significantly despite an apparent general increase in food choices in later components. Rates of responding were not affected systematically by VU0152099 treatment.
