## Supplemental Table S2 for "Effects of acute and repeated administration of the selective M_4_ PAM VU0152099 on cocaine vs. food choice in male rats"

**Supplemental Table S2.** Full ANOVA outcomes.

Table S2A. ANOVA outcomes for cocaine dose-response functions, repeated VU0152099 dosing

| | Cocaine dose | Treatment day | Interaction |
|---|---|---|---|
| % cocaine choice | F(1.9,13.6)=46.5, *p*<0.0001 | F(2.3,16.3)=4.1, *p*=0.03 | F(3.3,20.2)=1.7, *p*=0.19 |
| Baseline vs. | | | |
| Day 1 | F(1.3,9.3)=14.8, *p*=0.003 | F(1,7)=1.7, *p*=0.24 | F(1.5,10.8)=2.8, *p*=0.11 |
| Day 4 | F(1.8,12.8)=21.7, *p*<0.0001 | F(1,7)=0.59, *p*=0.47 | F(1.9,13.4)=0.92, *p*=0.42 |
| Day 5 | F(2.1,14.4)=35.9, *p*<0.0001 | F(1,7)=4.6, *p*=0.069 | F(1.9,13.6)=1.6, *p*=0.23 |
| Day 6 | F(1.6,11.2)=13.4, *p*=0.002 | F(1,7)=7.3, *p*=0.03 | F(1.3,7.6)=5.1, *p*=0.049 |
| Day 7 | F(2.0,13.9)=14.0, *p*=0.0005 | F(1,7)=25.0, *p*=0.002 | F(2.3,10.5)=10.3, *p*=0.003 |
| Post day 4 | F(2.3,16.7)=20.3, *p*<0.0001 | F(1,7)=1.9, *p*=0.21 | F(2.3,13.3)=2.7, *p*=0.10 |
| Cocaine reinforcers | F(1.6,11.3)=13.8, p=0.001 | F(3.3,23.0)=3.6, p=0.02 | F(20,125)=1.9, p=0.01 |
| Baseline vs. | | | |
| Day 1 | F(1.5,10.5)=40.7, *p*<0.0001 | F(1,7)=2.8, *p*=0.14 | F(1.6,11.2)=3.7, *p*=0.067 |
| Day 4 | F(1.8,12.7)=49.8, *p*<0.0001 | F(1,7)=2.0, *p*=0.21 | F(1.9,13.6)=1.1, *p*=0.37 |
| Day 5 | F(1.9,13.6)=95.0, *p*<0.0001 | F(1,7)=4.8, *p*=0.064 | F(1.8,12.3)=1.4, *p*=0.27 |
| Day 6 | F(1.7,11.6)=31.4, *p*<0.0001 | F(1,7)=3.5, *p*=0.11 | F(2.0,9.8)=2.3, *p*=0.15 |
| Day 7 | F(2.3,16.4)=20.3, *p*<0.0001 | F(1,7)=11.5, *p*=0.02 | F(3.2,14.3)=9.0, *p*=0.001 |
| Post day 4 | F(2.4,16.6)=57.3, *p*<0.0001 | F(1,7)=3.1, *p*=0.12 | F(2.3,12.4)=1.5, *p*=0.26 |
| Food reinforcers | F(2.6,18.0)=50.7, *p*<0.0001 | F(1.5,10.6)=1.0, *p*=0.37 | F(3.9,24.1)=2.2, *p*=0.10 |
| Total response rate | F(1.4,10.0)=4.6, *p*=0.048 | F(1.6,11.5)=0.78, *p*=0.46 | F(2.7,17.0)=1.2, *p*=0.34 |
| Cocaine+Food levers response rate | F(1.7,12.2)=2.2, *p*=0.16 | F(1.8,12.3)=1.3, *p*=0.30 | F(4.5,28.3)=0.58, *p*=0.70 |

Table S2B. ANOVA outcomes for session-wide cocaine and food intakes and cocaine $A_{50}$

| | Mixed-effect analysis | Test for trend |
|---|---|---|
| *Acute dosing, effect of* VU0152099 dose | | |
| Log(A50) cocaine dose | F(1.8,9.3)=3.9, *p*=0.06 | Nonlinear F(5,32)=3.8, *p*=0.008 |
| Cocaine intake / session | F(1.4,7.9)=3.5, *p*=0.09 | Nonlinear F(5,33)=3.0, *p*=0.02 |
| Food intake / session | F(1.6,8.9)=2.8, *p*=0.12 | Nonlinear F(5,33)=2.8, *p*=0.03 |
| *Repeated dosing, effect of day* | | |
| Log(A50) cocaine dose | F(2.6,14.9)=3.4, *p*=0.054 | Linear F(1,38)=11.3, *p*=0.002 |
| Cocaine intake / session | F(1.4,8.7)=2.5, *p*=0.15 | Linear F(1,32)=9.7, *p*=0.004 |
| Food intake / session | F(1.5,9.5)=1.0, *p*=0.37 | No significant trend, p>0.28 |

Table S2C. ANOVA outcomes for cocaine dose-response functions, acute VU0152099 dosing

| VU0152099 dose | Cocaine unit dose | Treatment day vs. baseline | Cocaine by treatment Interaction |
|---|---|---|---|
| *Percent cocaine choice* | | | |
| Vehicle | F(1.1,6.6)=47.6, *p*=0.0003 | F(1,6)=5.7, *p*=0.054 | F(1.3,7.0)=7.9, *p*=0.02 |
| 0.32 mg/kg | F(1.5,7.4)=28.6, *p*=0.0005 | F(1,5)=2.0, *p*=0.21 | F(1.5,7.3)=2.2, *p*=0.18 |
| 1.0 mg/kg | F(1.9,11.4)=27.8, *p*<0.0001 | F(1,6)=1.8, *p*=0.23 | F(1.6, 8.5)=2.0, *p*=41 |
| 1.8 mg/kg | F(1.3,7.8)=90.0, *p*<0.0001 | F(1,6)=5.0, *p*=0.068 | F(1.7,9.6)=2.3, *p*=0.15 |
| 3.2 mg/kg | F(2.2,13.4)=56.7, *p*<0.0001 | F(1,6)=2.1, *p*=0.20 | F(1.7,10.5)=0.75, *p*=0.48 |
| 5.6 mg/kg | F(1.7,6.8)=50.0, *p*<0.0001 | F(1,4)=1.7, *p*=0.35 | F(1.4,5.3)=0.88, *p*=0.35 |
| *Cocaine reinforcers taken* | | | |
| Vehicle | F(1.2,7.0)=29.0, *p*=0.0008 | F(1,6)=6.7, *p*=0.04 | F(1.5,8.7)=7.5, *p*=0.02 |
| 0.32 mg/kg | F(1.4,7.4)=11.6, *p*=0.007 | F(1,5)=3.1, *p*=0.14 | F(1.7,8.5)=2.3, *p*=0.16 |
| 1.0 mg/kg | F(1.5,9.3)=18.3, *p*=0.0009 | F(1,6)=2.3, *p*=0.18 | F(1.9,11.5)=2.4, *p*=0.13 |
| 1.8 mg/kg | F(1.3,7.6)=42.3, *p*=0.0002 | F(1,6)=8.5, *p*=0.03 | F(1.7,10.0)=2.2, *p*=0.16 |
| 3.2 mg/kg | F(2.0,11.8)=31.1, *p*<0.0001 | F(1,6)=3.8, *p*=0.10 | F(1.4,8.7)=1.8, *p*=0.22 |
| 5.6 mg/kg | F(1.6,6.4)=31.7, *p*=0.0006 | F(1,4)=1.0, *p*=0.37 | F(1.8,7.3)=1.2, *p*=0.35 |
| *Liquid food reinforcers taken* | | | |
| Vehicle | F(1.1,6.5)=45.3, *p*=0.0003 | F(1,6)=9.1, *p*=0.02 | F(1.4,8.0)=7.6, *p*=0.02 |
| 0.32 mg/kg | F(1.9,9.6)=15.3, *p*=0.001 | F(1,5)=3.3, *p*=0.13 | F(1.4,6.9)=1.8, *p*=0.23 |
| 1.0 mg/kg | F(1.9,11.5)=31.1, *p*<0.0001 | F(1,6)=0.12, *p*=0.74 | F(1.5,9.2)=2.3, *p*=0.16 |
| 1.8 mg/kg | F(2.5,15.2)=43.8, *p*<0.0001 | F(1,6)=0.97, *p*=0.36 | F(2.1,12.4)=2.8, *p*=0.10 |
| 3.2 mg/kg | F(2.0,12.2)=49.1, *p*<0.0001 | F(1,6)=0.50, *p*=0.51 | F(1.7,10.6)=1.7, *p*=0.23 |
| 5.6 mg/kg | F(1.7,7.1)=43.9, *p*<0.0001 | F(1,4)=1.8, *p*=0.26 | F(1.3,5.3)=0.74, *p*=0.47 |
| *Responses per minute, all levers* | | | |
| Vehicle | F(1.6,9.8)=9.7, *p*=0.006 | F(1,6)=2.9, *p*=0.14 | F(1.9,11.2)=0.74, *p*=0.49 |
| 0.32 mg/kg | F(1.4,8.4)=5.4, *p*=0.04 | F(1,6)=0.63, *p*=0.46 | F(1.4, 8.9)=2.9, *p*=0.12 |
| 1.0 mg/kg | F(1.3,7.9)=11.8, *p*=0.007 | F(1,6)=0.04, *p*=0.85 | F(1.8,10.6)=0.78, *p*=0.47 |
| 1.8 mg/kg | F(1.3,8.1)=14.9, *p*<0.003 | F(1,6)=0.15, *p*=0.71 | F(1.5,8.9)=1.2, *p*=0.34 |
| 3.2 mg/kg | F(1.3,8.1)=14.9, *p*=0.003 | F(1,6)=0.15, *p*=0.71 | F(1.5,8.9)=1.2, *p*=0.34 |
| 5.6 mg/kg | F(1.3,5.1)=11.3, *p*=0.02 | F(1,4)=0.82, *p*=0.41 | F(2.4,9.7)=3.9, *p*=0.055 |
| *Responses per minute, reinforcer selection levers* | | | |
| Vehicle | F(1.8,10.7)=1.2, *p*=0.35 | F(1,6)=1.3, *p*=0.29 | F(2.7,16.0)=1.4, *p*=0.27 |
| 0.32 mg/kg | F(3.2,15.9)=1.5, *p*=0.24 | F(1,5)=41.6, *p*=0.001 | F(3.0,14.8)=0.74, *p*=0.54 |
| 1.0 mg/kg | F(1.2,7.2)=1.6, *p*=0.25 | F(1,6)=0.60, *p*=0.47 | F(1.4,8.8)=1.1, *p*=0.36 |
| 1.8 mg/kg | F(2.1,12.9)=3.4, *p*=0.064 | F(1,6)=1.2, *p*=0.32 | F(2.0,11.8)=1.3, *p*=0.30 |
| 3.2 mg/kg | F(2.4,14.3)=2.8, *p*=0.09 | F(1,6)=0.21, *p*=0.66 | F(1.9,11.3)=0.99, *p*=0.40 |
| 5.6 mg/kg | F(2.3,9.1)=2.0, *p*=0.19 | F(1,4)=14.0, *p*=0.02 | F(2.0,8.2)=0.52, *p*=0.52 |

Table S2D. ANOVA outcomes for cocaine dose-response functions with VU0152100 in CalDAG-GEFI knockout mice

| | 3-way ANOVA |
|---|---|
| Pretreatment | F(1,42)=19.4, p=0.0001 |
| Cocaine dose | F(3,42)=8.08, p=0.004 |
| CalDAG-GEFI genotype | F(1,42)=12.6, p=0.001 |
| Cocaine by pretreatment interaction | F(3,42)=4.28, p=0.02 |
| Cocaine by genotype interaction | F(3,42)=3.35, p=0.07 |
| Pretreatment by genotype interaction | F(1,42)=0.25, p=0.62 |
| 3-way interaction | F(3,42)=0.57, p=0.58 |
