## Supplemental Figure S3 for "Effects of acute and repeated administration of the selective M_4_ PAM VU0152099 on cocaine vs. food choice in male rats"

**Supplemental Figure S3.** Percent cocaine choice in individual rats at baseline and days 5 and 7 of repeated VU0152099 treatment.

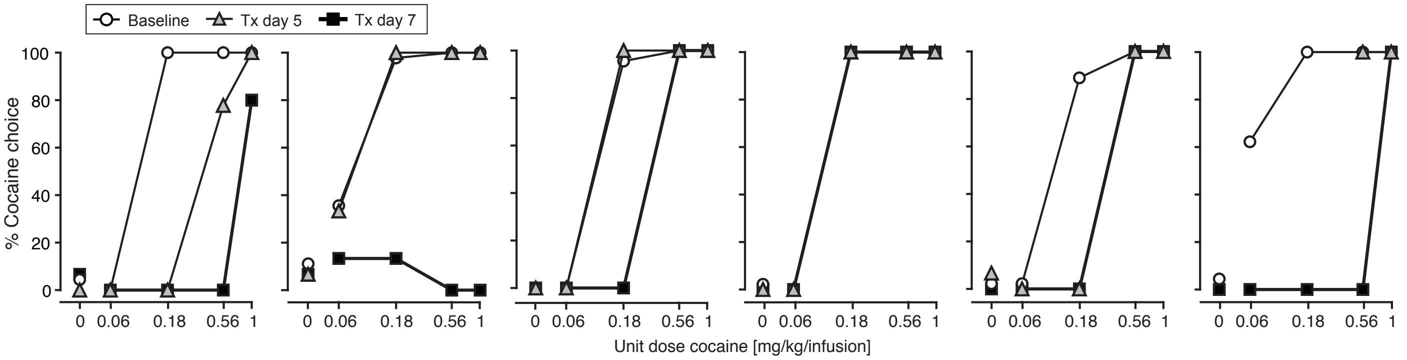


Five of six rats showed some shift in behavior allocation away from cocaine taking towards food taking by day seven, with more varied response earlier in the treatment.
