## Supplemental Figure S4 for "Effects of acute and repeated administration of the selective M_4_ PAM VU0152099 on cocaine vs. food choice in male rats"

**Supplemental Figure S4.** Repeated vehicle treatment

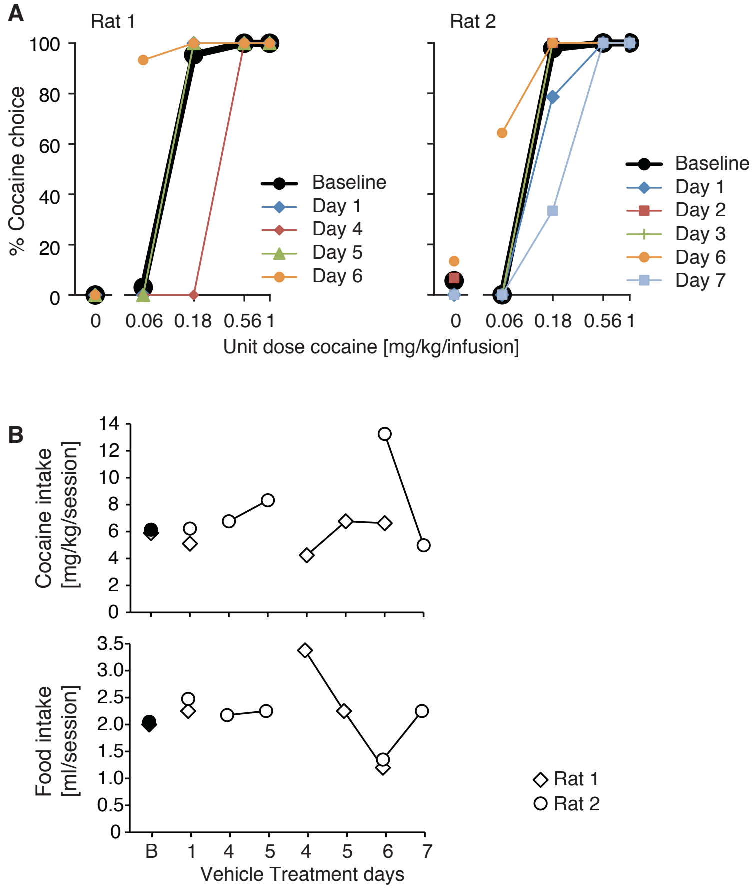


(A) Percent cocaine choice allocation in two rats that received repeated vehicle injections, all days tested; rat 1 started on a Friday, rat 2, on a Wednesday. (B) Session-wide cocaine and food intakes in each rat as a function of successive days (baseline levels shown as filled symbols, treatment days as open symbols; baseline level symbols overlap).
