## Supplemental Figure S5 for "Effects of acute and repeated administration of the selective M_4_ PAM VU0152099 on cocaine vs. food choice in male rats"

**Supplemental Figure S5.** Rates of responding, acute VU0152099 dosing

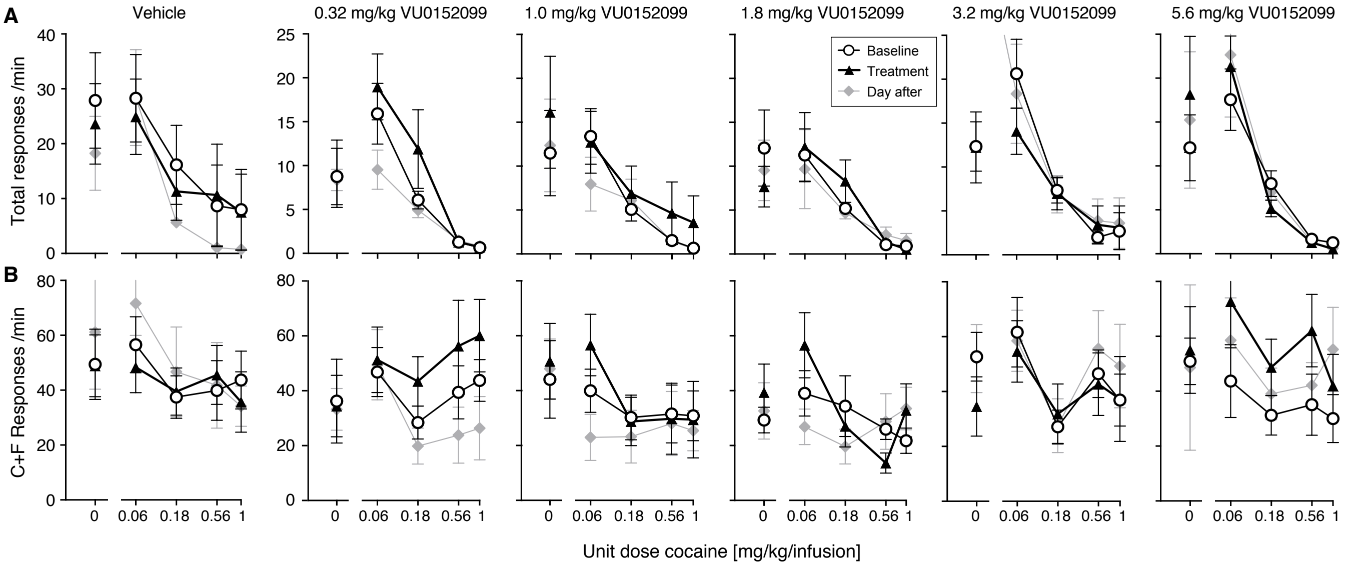


Total rates (A, all three levers), and rate of responding on the two reinforcer-selection levers (B) as functions of cocaine unit dose, for each acute dose of VU0152099. Total rates but not reinforcer-selection lever rates were related to cocaine unit dose, indicating that the determining factor in the rate dependency was time between reinforcer selection (and thus reinforcer delivery) and starting the next chain by pressing the observer lever. This is consistent with our previous findings using this assay. VU0152099 treatment did not decrease rates of responding at any of the doses tested, but increased rates of responding on the reinforcer-selection levers at 0.32 and 5.6 mg/kg (see Table S2C for statistical outcomes).
