## Supplemental Figure S6 for "Effects of acute and repeated administration of the selective M_4_ PAM VU0152099 on cocaine vs. food choice in male rats"

**Supplemental Figure S6.** Session-summary measures, acute VU0152099 dosing

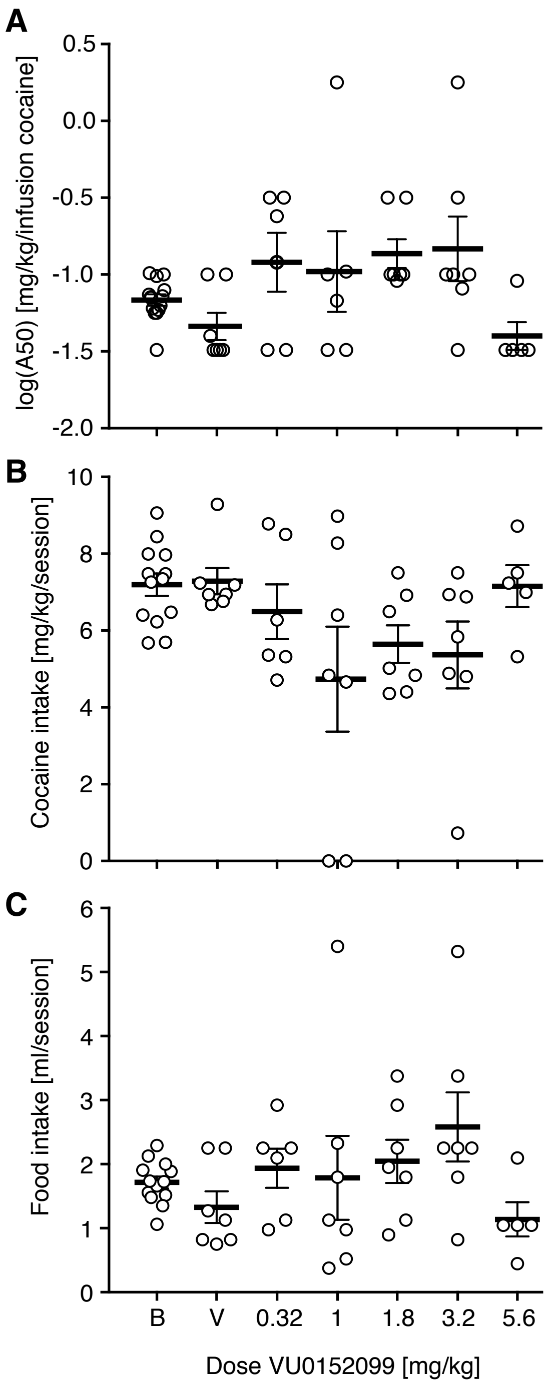


Session-wide cocaine choice $A_{50}$ (A) and cocaine (B) and food (C) intakes as functions of acute VU0152099 dose. “B”: baseline performance, “V”: vehicle. Data represent individual rats as well as group means with s.e.m.
